## Supplementary figures and images for "Alterations in protein translation and carboxylic acid catabolic processes in diabetic kidney disease"

### Supplemental Figure 1

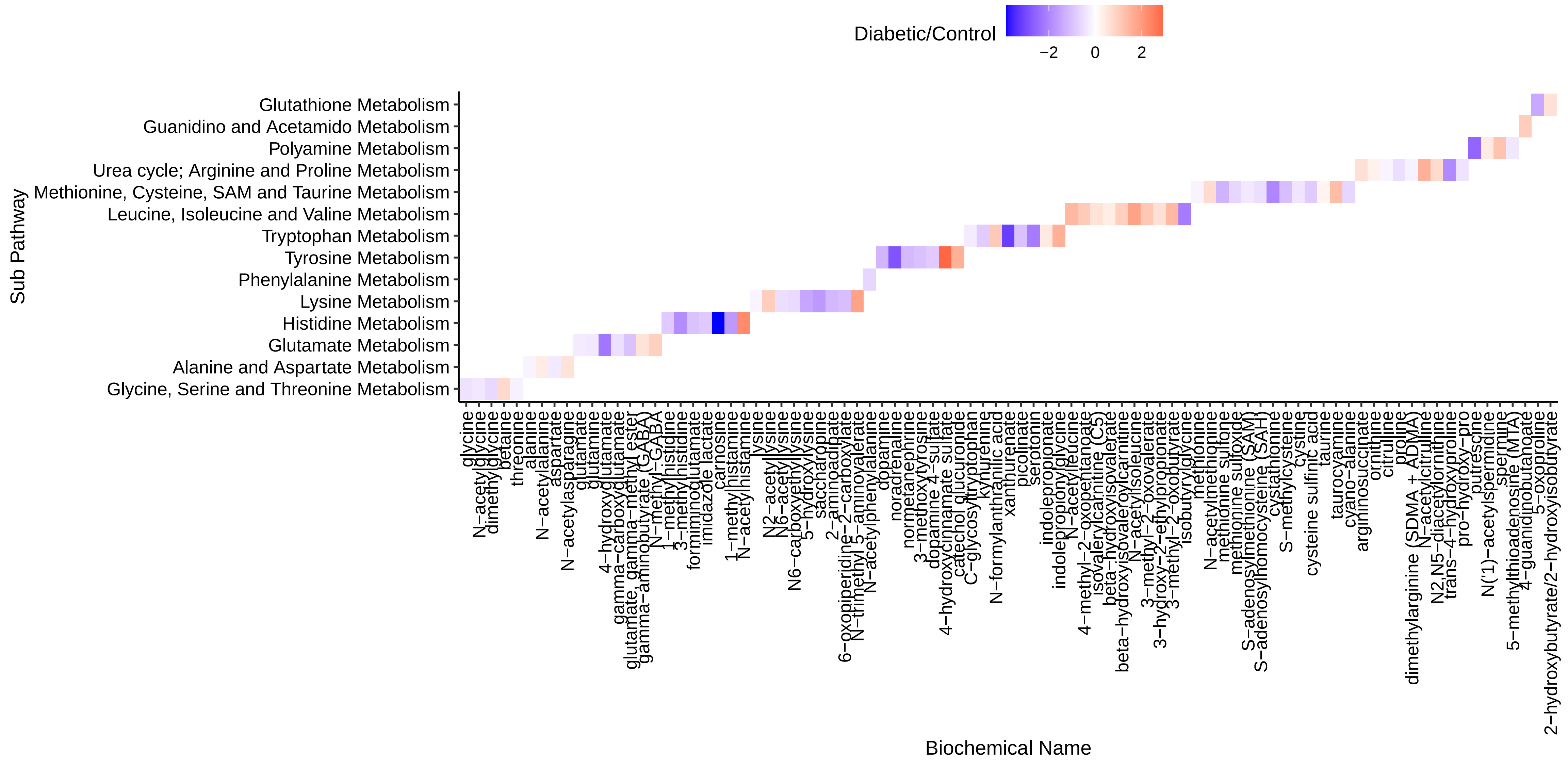
